# Comparative Genomics Reveals Deep Conservation and Lineage-specific Evolution of Mammalian microRNAs

**DOI:** 10.64898/2025.12.23.696236

**Authors:** Sarahjane Power, Zixia Huang

## Abstract

microRNAs (miRNAs) are central post-transcriptional regulators that shape cellular functions, development, and phenotypic diversity. Yet their evolutionary history in mammals remains poorly resolved, largely due to incomplete genome-wide annotation and the difficulties in establishing miRNA orthology. Using a homology-based framework paired with genome synteny for orthology assignment, we characterised miRNA families and investigated their evolutionary dynamics across mammals. Compared to Infernal and MirMachine using a consensus pseudo-ground-truth framework, our method demonstrated better sensitivity-specificity balance, allowing divergent miRNA detection while reducing false positives. Analysis of 73 mammalian genomes revealed major miRNA family gains at the ancestral nodes of Eutheria, Marsupialia, and Haplorrhini, contrasted with substantial losses in Eulipotyphla and Scandentia. Copy number evolution was largely conserved across families, with only a few families (e.g., MIR-145, MIR-2285) showing lineage-specific duplication acceleration. Comparative genome microsynteny further resolved orthology for the multi-copy family LET-7, spanning eight deeply conserved syntenic blocks, as well as for 143 individual miRNAs. We showed that most miRNAs are deeply conserved, with the seed region being most constrained, followed by the mature, star, and loop regions. Within the seed region, positions 4-6 displayed higher conservation than other seed positions, consistent with their roles in core target recognition. Ancestral seed-state reconstruction further revealed diverse evolutionary signatures, including ancestral shifts, recurrent substitutions, and homoplasious changes. Collectively, these findings provide an unprecedented comparative overview of mammalian miRNA evolution and create new opportunities to investigate how shifts in miRNA repertoires and sequence features may underlie lineage-specific regulatory and phenotypic diversification.

## Introduction

MicroRNAs (miRNAs) are small, non-coding RNAs that function as central post-transcriptional regulators across metazoans, fine-tuning gene expression programs with remarkable precision (Bartel 2004; Filipowicz et al. 2008). Mature miRNAs are produced through a conserved biogenesis pathway in which primary miRNA transcripts (pri-miRNAs) are sequentially processed by the Drosha-DGCR8 complex in the nucleus and Dicer in the cytoplasm to yield a ∼22-nt duplex, from which one strand is selectively loaded onto the RNA-induced silencing complex (RISC). Acting primarily through base-pairing between their 5’ seed region and complementary sites within target mRNAs, miRNAs guide RISC to specific transcripts, resulting in translational repression or mRNA destabilisation. Since their discovery, miRNAs have been demonstrated to perform essential roles, including development timing (Pasquinelli and Ruvkun 2002), cell differentiation (Ivey and Srivastava 2010; Shenoy and Blelloch 2014), and physiological homeostasis (Dumortier et al. 2013), with dysregulation implicated in a broad range of diseases (Peng and Croce 2016; Kapplingattu et al. 2025). Owing to their pervasive impact, considerable efforts have been devoted to elucidating the full scope of miRNA mechanisms (Treiber et al. 2019; Diener et al. 2024), their roles in shaping phenotypic complexity (Heimberg et al. 2008; Zolotarov et al. 2022), and their emerging potential as therapeutic targets (Chakraborty et al. 2017; Diener et al. 2022).

miRNA innovation has long been associated with increases in organismal complexity, with major waves of emergence coinciding with key evolutionary transitions, such as bilaterians, vertebrates, and placental mammals (Berezikov 2011; Wang et al. 2024). Several deeply conserved miRNA families, including LET-7 and MIR-9, arose early in metazoan evolution and became embedded in core developmental gene regulatory networks that govern processes such as cell fate specification, neurogenesis, and axis patterning (Coolen et al. 2012; Papaioannou et al. 2013; Madelaine et al. 2017). In contrast, some miRNAs display far more dynamic evolutionary trajectories. For example, the gain of MIR-196 in vertebrates introduced a new regulatory layer to *Hox* gene patterning, directly repressing HOX transcripts and reshaping limb and axial skeletal morphology (He et al. 2011). Another striking illustration comes from recent work in butterflies showing that MIR-193 is the key regulator of melanic wing patterns, as its knockout in multiple lepidopteran lineages resulted in the loss of dark pigmentation (Tian et al. 2024). Likewise, the gain of MIR-279 in insects proved essential for the formation of the CO_2_-sensing neural circuit, with knockout experiments in fruit flies demonstrating that loss of this single miRNA abolishes CO_2_ avoidance behaviour (Hartl et al. 2011). Beyond miRNA gains and losses, alternation of seed regions, the primary determinant of target specificity, can also introduce new phenotypes. One of the most well-known examples is a single base substitution in the MIR-96 seed region that causes dominant progressive hearing loss in mice (Zhu et al. 2024), while a similar seed change in MIR-204 leads to autosomal dominant retinal dystrophy in medaka fish (Conte et al. 2015). These examples highlight that both emergence of new miRNAs and subtle modifications within existing miRNAs can contribute to phenotypic diversification across the tree of life.

Although these studies reveal the profound effects a single miRNA family can have on phenotype, they offer limited insight into how the broader miRNA repertoire naturally evolves or how such evolutionary changes contribute to organismal diversity. Recent genome-wide miRNA studies have primarily focused on improving miRNA annotation, and characterising patterns of gain, loss, and duplication across lineages, as well as exploring how these evolutionary dynamics may relate to phenotypic traits. This includes advances in covariate-model-based miRNA annotation frameworks (Umu et al. 2023; Clarke et al. 2025; Vanek et al. 2025), analyses of miRNA trajectories following whole-genome duplication events (Desvignes et al. 2021; Peterson et al. 2022), investigations into the origins and expression dynamics of miRNAs in birds and mammals (Fromm et al. 2013; Meunier et al. 2013; Penso-Dolfin et al. 2018; Taylor et al. 2023), and large-scale survey of miRNA target evolution (Xu et al. 2013; Simkin et al. 2020). However, far less is known about how miRNA families evolve at the level of copy number and sequence, features with direct consequences for regulatory function but rarely explored beyond gain and loss patterns. To narrow these gaps, we focus here on the evolution of miRNA copy number and sequence across mammals, a lineage with exceptional ecological and phenotypic diversity that remains underexplored from an evolutionary miRNA perspective.

The difficulty stems from the challenges of obtaining comprehensive genome-wide annotations that capture both conserved and lineage-specific families across species. Most current miRNA identification tools rely on small RNA-Seq data (Friedlander et al. 2012; Vitsios et al. 2017; Aparicio-Puerta et al. 2022), which can detect context-specific and lineage-specific families but are constrained by sample availability and tissue representation, often resulting in incomplete or biased miRNA catalogs. To address these limitations, genome-based approaches such as Infernal and MirMachine have been developed. Infernal uses Rfam covariate models (CMs) and can, in principle, detect a wide range of noncoding RNAs, including miRNAs, offering flexibility and applicability across diverse species (Nawrocki and Eddy 2013). However, many Rfam CMs were built from limited or outdated sequence sets, leading to poorly defined models and high false-positive rates (Fromm et al. 2015). MirMachine, in contrast, relies on the rigorously curated MirGeneDB CMs, enabling far more accurate and specific annotation of conserved miRNA families and substantially reducing false positives (Umu et al. 2023). Yet this strength also comes with a major limitation because its models are built primarily around deeply conserved families. As a result, MirMachine preferentially recovers ancient miRNAs while systematically missing many validated but lineage-specific miRNAs represented in MirGeneDB, which are precisely the families most informative for understanding the evolutionary diversification of miRNAs. Adding to this challenge, resolving miRNA evolution also requires establishing reliable orthology assignment, a task complicated by the short length of miRNA precursors, their limited sequence conservation outside the seed region, and the frequent duplication or loss of miRNA genes across lineages, all of which obscure clear one-to-one evolutionary relationships (Fromm et al. 2015).

Thus, a fundamental question remains: How do miRNA families evolve in both copy number and sequence, particularly in their seed regions, across mammals? To address this question, we developed a homology-based framework that surveyed well-curated MirGeneDB miRNA families across mammalian genomes and integrated genome microsynteny to generate robust and biologically informed orthology assignments. We first compared our approach with Infernal and MirMachine on ten representative mammalian genomes using a consensus pseudo-ground-truth framework, demonstrating its effectiveness relative to existing tools. We then applied this approach to annotate miRNA families across 73 mammalian genomes, infer their gain and loss events across the mammalian phylogeny, and characterise patterns of their copy number evolution. Through genome microsynteny analyses, we resolved orthology relationships for miRNA families, including the complex LET-7 family. We further examined the conservation of distinct precursor features, focusing in particular on the evolutionary dynamics of seed regions. Finally, by reconstructing ancestral seed states we uncovered a range of evolutionary signatures, including ancestral shifts, recurrent substitutions, and homoplasious changes, that collectively reveal how miRNA seed sequences have diversified during mammalian evolution. Together, this study provides one of the most comprehensive evolutionary portraits of mammalian miRNAs, offering a foundation for future work investigating how changes in miRNA repertoires and sequences may have contributed to lineage-specific regulatory and phenotypic diversity.

## Results

### Performance comparisons of three in silico miRNA identification pipelines

We evaluated the performance of our homology-based miRNA prediction method against two established pipelines, Infernal and MirMachine. To ensure a fair comparison, each method was evaluated using its own predefined set of mammalian miRNA families (see Methods). The composition of these query sets varied markedly, with only 228 families (21.8%) shared across all methods (Fig. 1A). Infernal contained the most unique families (N = 454, 43.4%), followed by the homology-based method (N = 273, 26.1%). Many of the Infernal-specific CMs were constructed from significantly fewer input sequences than those in the shared set (*P* < 2.2 × 10^-16^, Mann-Whitney *U* test; Fig. S1A). The families uniquely predicted by the homology-based method correspond to clade- or species-specific miRNA families documented in MirGeneDB but absent from the MirMachine query set.

**Figure 1:**
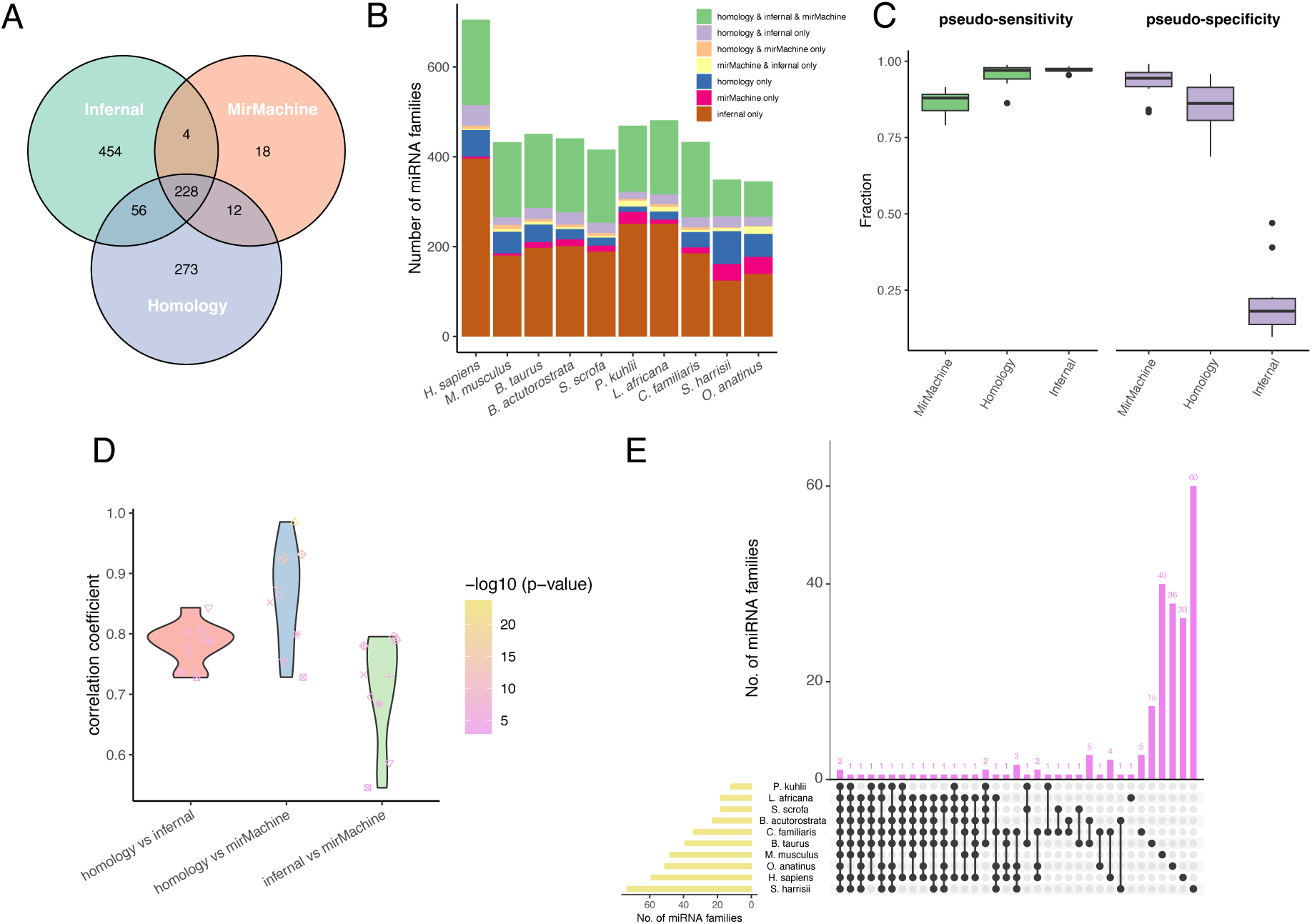
Comparison of miRNA identification results across three pipelines (the homology-based approach, Infernal, and MirMachine). **A)** Number of miRNA families used as query sets for *in silico* identification in each pipeline, and their overlap. **B)** Number of miRNA families predicted across ten representative mammalian genomes by each pipeline, and their overlap. **C)** Evaluation of the pseudo-sensitivity and pseudo-specificity of three prediction methods. Values reflect pseudo-sensitivity and pseudo-specificity calculated using a consensus pseudo-ground-truth framework (See Methods). **D)** Pairwise comparison of Spearman correlation coefficients based on miRNA family copy numbers jointly identified by all three pipelines. **E)** UpSetR plot showing the overlap of miRNA families uniquely predicted by our homology-based pipeline across ten mammalian genomes.

Across the ten species, the homology-based method recovered an average of 216.2 ± 41.3 miRNA families per genome, compared with 179.2 ± 25.7 predicted by MirMachine, and 392.1 ± 109.2 predicted by Infernal. *Homo sapiens* showed the highest number of predicted families (N = 705), whereas *Ornithorhynchus anatinus* exhibited the lowest (N = 345) (Fig. 1B). When partitioning predictions by method, the largest categories were miRNA families predicted exclusively by Infernal (45.7% ± 6.5%) and those detected concurrently by all three methods (33.0% ± 6.4%) (Fig. S1B). The complete overlap of miRNA families predicted by three approaches across ten species is shown in Fig. S1C.

Using the consensus-based pseudo-ground-truth framework (See Methods), clear differences emerged among the three approaches. Infernal achieved the highest pseudo-sensitivity (97.2% ± 0.8%), with the homology-based approach performing similarly (95.4% ± 3.7%) and MirMachine detecting fewer consensus families (86.7% ± 4.2%) (Fig. 1C). The pattern reversed when examining pseudo-specificity. MirMachine performed best (93.0% ± 5.4%), followed by the homology-based approach (85.4% ± 8.3%), whereas Infernal showed markedly lower pseudo-specificity (21.6% ± 12.1%) due to its larger number of unique predictions (Fig. 1C).

For miRNA families detected by all three methods, we assessed the concordance of copy-number estimates across methods. All pairwise Spearman correlations were positive (ρ > 0) and statistically significant (FDR < 0.05). Concordance was highest between the homology-based pipeline and MirMachine (ρ = 0.864 ± 0.083), lowest between Infernal and MirMachine (ρ = 0.713 ± 0.087), and intermediate between the homology-based method and Infernal (ρ = 0.785 ± 0.034) (Fig. 1D). Examination of families unique to the homology-based method identified 40 (17.4%) miRNA families present in more than one species, including two detected in all ten species (Fig. 1E).

### Genome-wide miRNA family prediction in 73 mammalian species using a homology-based method

To characterise miRNA evolutionary patterns across mammals, we predicted 568 miRNA families across 73 species using the homology-based method. On average, 224.3 ± 42.2 families were predicted per species, with *Pongo pygmaeus* showing the highest number (N = 303) and *Sorex araneus* the lowest (N = 151) (Figs. 2A-B). Among mammalian orders, Primates exhibited the highest mean number of predicted families (278.6 ± 28.9). Notably, 54 of the 73 species currently lack curated miRNA annotations in MirGeneDB, and for the 19 species with existing entries, our homology-based method increased family counts by 10.7% ± 5.5% on average (Fig. 2B). Across all species, an average of 155.7 ± 34.9 miRNA families were present as single-copy families, 62.7 ± 7.3 had two to ten copies, and only 5.7 ± 2.6 had more than ten copies (Figs. S2A-B).

**Figure 2:**
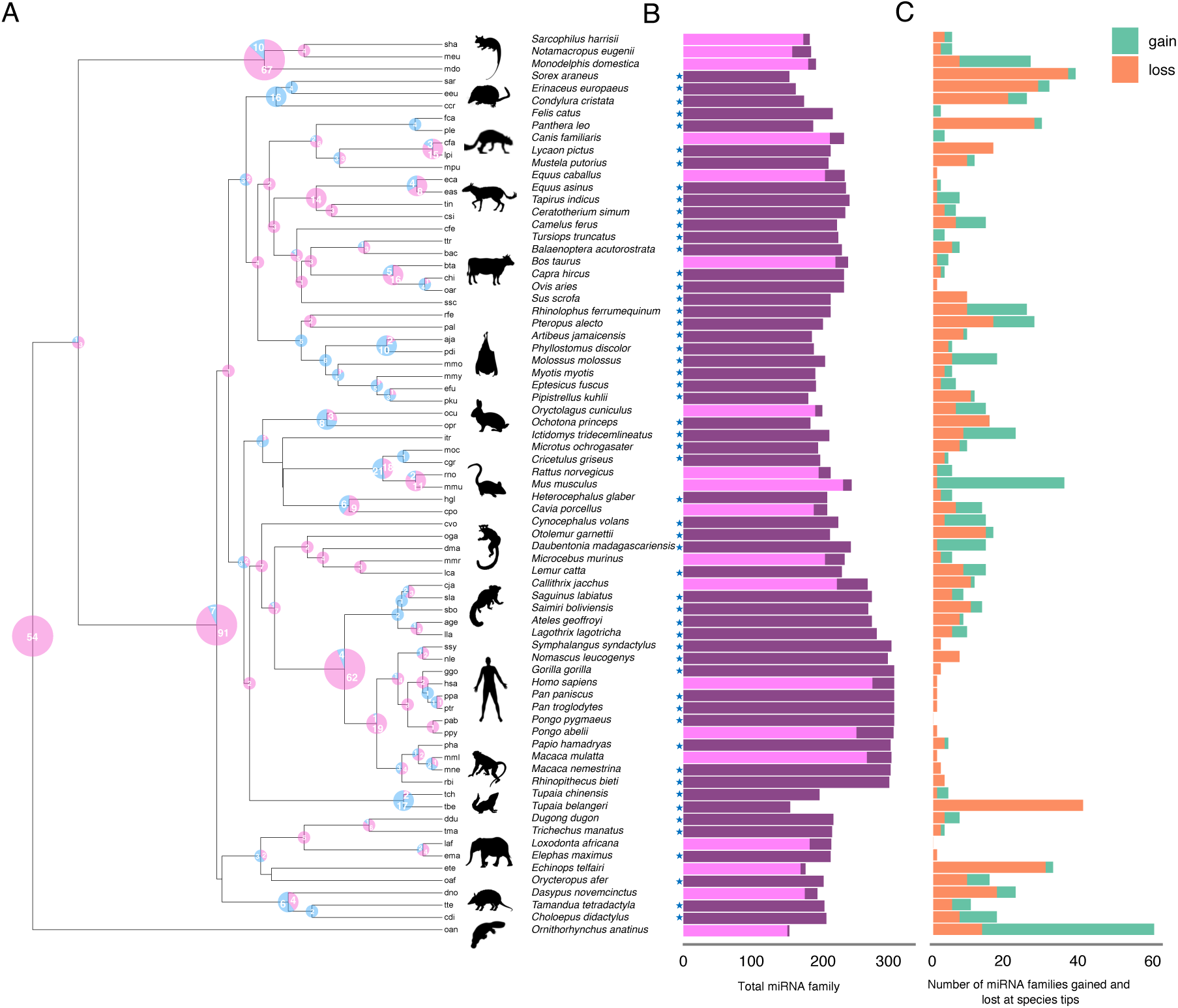
miRNA family prediction across 73 mammalian genomes using the homology-based pipeline. **A)** Inferred miRNA family gains and losses mapped onto the mammalian phylogeny. The phylogenetic tree was obtained from TimeTree v5. Numbers on internal nodes represent the number of gains (pink) and losses (blue) estimated using maximum-likelihood ancestral state reconstruction. Node size reflects the total number of evolutionary events (gains + losses): small (0 – 50), medium (50 – 100), and large (>100). **B)** Number of miRNA families predicted per species. Light purple bars represent families curated in MirGeneDB; dark purple bars represent additional families identified by our homology-based pipeline. An asterisk (*) marks species that currently lack miRNA annotations in MirGeneDB. **C)** Number of miRNA family gains and losses inferred for each species (tree tips).

### miRNA family gains and losses across the mammalian phylogeny

Next, we inferred miRNA family gains and losses along the established mammalian phylogeny using maximum-likelihood ancestral state reconstruction (See Methods). The largest numbers of inferred gains were observed at the Eutherian (91 gains), Marsupial (67 gains), and Haplorrhine (62 gains) nodes, whereas the highest numbers of inferred losses occurred in Scandentia (17 losses) and Eulipotyphla (16 losses) (Fig. 2A). At the species level, *O. anatinus* exhibited the highest number of family gains (46), followed by *Mus musculus* (34), while the highest number of family losses was detected in *Tupaia belangeri* (40), followed by *S. araneus* (36) (Fig. 2C). Among all families that experienced losses, approximately two thirds showed only a few events, with 139 (66.5%) families being independently lost no more than three times across the phylogeny (Fig. S2C). In contrast, a small number of miRNA families (e.g. MIR-12472, MIR-384, and MIR-3593) exhibited more than 10 independent losses (Fig. S2C).

### Evolutionary analyses of miRNA family copy numbers

We performed a principal component analysis (PCA) using copy number estimates for 568 miRNA families across 73 species. The first two components separated species into three major clusters: non-eutherian mammals (monotremes and marsupials), primates and other eutherian species (Fig. 3A). To further explore miRNA copy number evolution, we estimated the evolutionary signal (Pagel’s λ) and the evolutionary rate (σ²) for each family (See Methods). Most miRNA families showed strong phylogenetic signal (high λ) and slow copy number evolution (low σ²), suggesting that copy numbers are largely conserved among closely related species (Fig. 3B, Fig. S3). A smaller subset showed accelerated turnover (high σ²), including MIR-145, MIR-2285 and MIR-544, which displayed markedly higher copy numbers in specific clades (Fig. 3C). For example, MIR-2285 exhibited considerably higher copy numbers in species belonging to Cetartiodactyla (Fig. 3C). However, many of these copies overlapped low-complexity regions or annotated transposable elements (TEs) across species except for MIR-506, suggesting possible functional loss (Fig. 3D).

**Figure 3:**
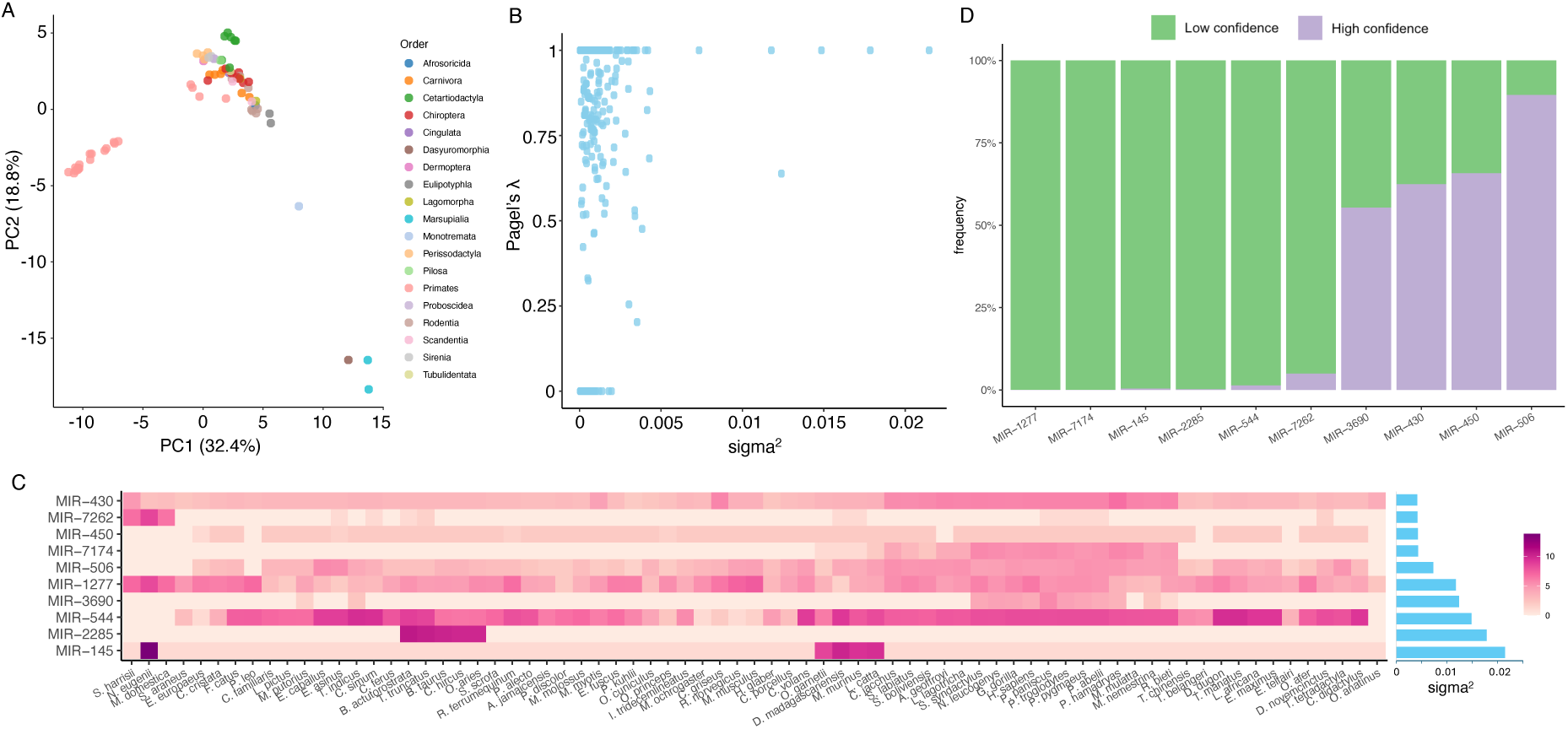
Copy number variation of miRNA families across mammals. **A)** Principal component analysis (PCA) based on copy numbers of 568 miRNA families across 73 mammalian genomes. **B)** Relationship between phylogenetic signals (Pagel’s λ) and evolutionary rate (σ²) across the same 568 miRNA families. Each point represents a miRNA family. Pagel’s λ reflects how strong copy number variation follows the phylogeny, and σ² represents the estimated rate of copy number evolution under a Brownian motion model. **C)** Heatmap showing copy numbers (log-transformed) of the top 10 most variable miRNA families (ranked by σ²) across 73 species. **D)** Frequency of high- and low-confidence miRNA copies within the top 10 most variable miRNA families. Low-confidence copies correspond to precursor sequences overlapping annotated transposable elements.

### Syntenic analyses of LET-7 family members

The short length and high sequence conservation of miRNAs often confound orthology assignment, especially for large, multi-copy families. To address this limitation, we combined comparative genome synteny analyses with our homology-based predictions (see Methods). Using this integrated approach, we resolved orthology relationships among members of LET-7 family, an evolutionarily ancient, functionally important, and multi-copy family comprising 9 – 14 paralogs across mammalian genomes.

We identified eight deeply conserved synteny blocks containing 12 copies of LET-7 across mammalian species (Fig. 4; Fig. S4). Three of these paralogs (Let-7-P2a1, Let-7-P2b1, Let-7-P2c1) occurred as a tandem cluster flanked by *PTPDC1* and *ZNF169* in most species examined (synteny block 1; Fig. 4). Two paralogs (Let-7-P2a3, Let-7-P2b3) were located within the intron of *HUWE1* in most species, except in *Choloepus didactylus*, where they were positioned between *VSIG4* and *SMC1A*, and in *Panthera leo*, where both paralogs were absent (synteny block 3; Fig. 4). Interestingly, the two marsupial species (*Sarcophilus harrisii* and *Monodelphis domestica*) appeared to retain only a single copy of this paralog pair (synteny block 3; Fig. 4). Similarly, another paralog pair (Let-7-P2a2, Let-7-P2b2) was located between *WNT7B* and *PPARA* in most species, whereas *S. harrisii* retained only one copy, *Pteropus alecto* contained three copies, and *Condylura cristata* lacked both copies entirely (synteny block 4; Fig. S4). The remaining five synteny blocks each contained a single Let-7 paralog, with evidence of species- or clade-specific losses or microsynteny rearrangements across the phylogeny (Fig. 4; Fig. S4).

**Figure 4:**
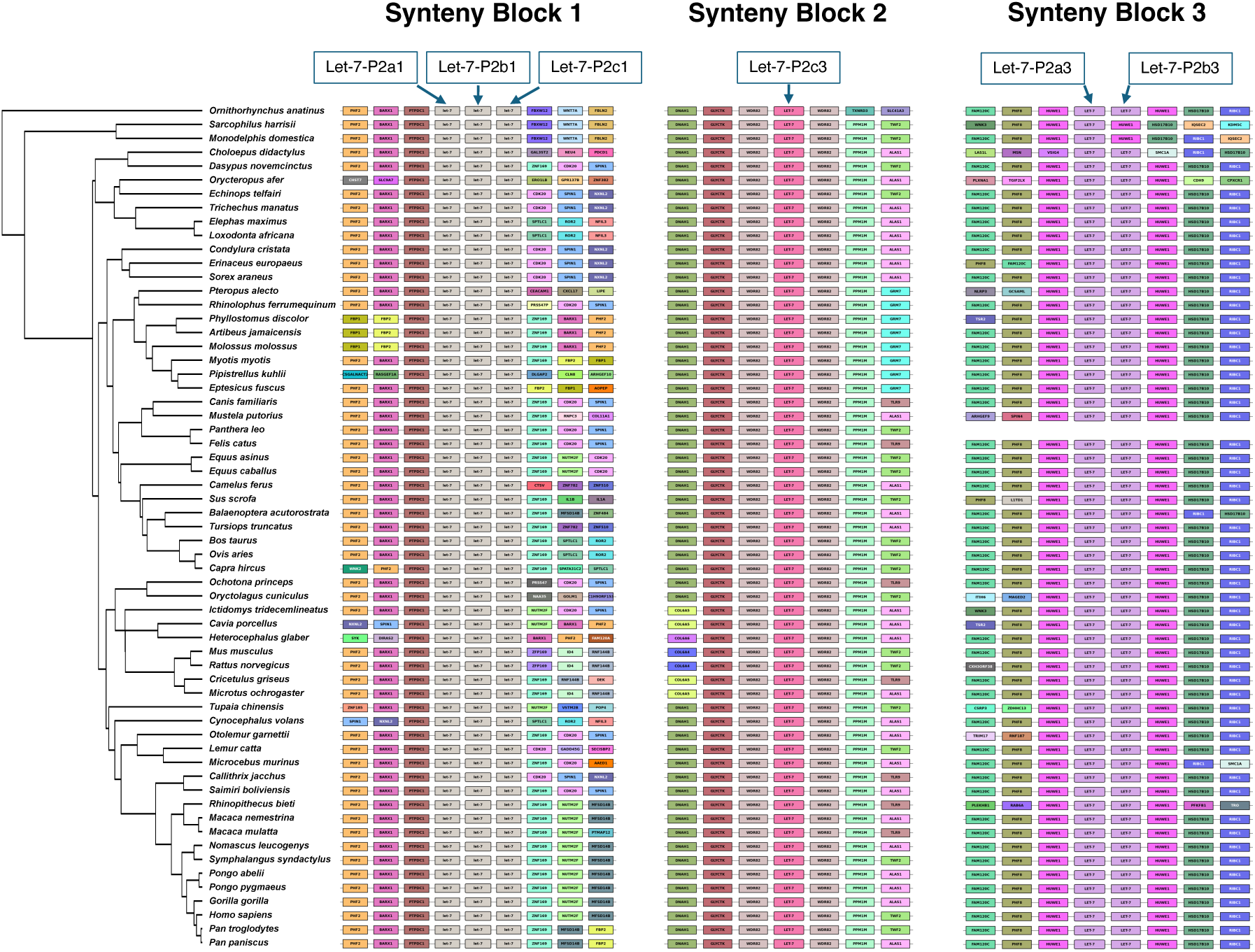
Comparative genome microsynteny illustrating orthology relationships among multi-copy LET-7 family members. The left panel shows the phylogeny of the 61 species included in the synteny analysis, with the tree obtained from TimeTree v5. The right panels depict three conserved syntenic blocks that contain LET-7 family members. Within each block, LET-7 loci are positioned centrally, with three upstream and three downstream protein-coding genes displayed. Colors represent different protein-coding genes and are consistent across species. The empty line indicates the species in which corresponding LET-7 members are absent. Five additional conserved syntenic blocks that contain LET-7 members are shown in Fig. S4.

In contrast, we also identified genomic regions that harbored clade- or species-specific Let-7 paralogs. For example, *C. didactylus* was predicted to contain 16 copies, the highest number among all species examined. In addition to the 12 members located within the eight conserved synteny blocks, this species possessed an additional tandem cluster of four paralogs situated between *KDM5C* and *TSR2* (Fig. S5). Furthermore, *Lemur catta*, *Symphalangus syndactylus*, and *Nomascus leucogenys* each possessed a single Let-7 paralog between *TMEM132E* and *CCL1* (or *TMEM132E* and *ASIC2* in *L. catta*), indicating a clade-specific duplication event (Fig. S5). Although these clade-specific paralogs harbored seed sequences that diverged from the canonical Let-7 seed, none were associated with TEs or low-complexity regions.

### Analyses of orthologous miRNAs

Using this approach, we also resolved and confirmed orthology relationships for 127 miRNA families, including 111 single-copy families and 16 two-copy families, totaling 143 individual precursor miRNAs. These orthologous miRNAs were embedded in deeply conserved synteny blocks, as indicated by the high recurrence of the five most frequently shared protein-coding genes, providing strong support for the robustness of our orthology assignments (Fig. 5A; see Methods). Based on these synteny-confirmed orthologs, we generated multiple sequence alignments for each orthologous miRNA across species. According to human annotations in MirGeneDB, these 143 precursor miRNAs corresponded to 152 mature regions (including cases where both mature and co-mature regions were defined), 131 star regions (with some lacking annotated star sequences), and 142 loop regions (with miR-451 lacking a defined loop region). Each precursor miRNA was represented by sequence data from at least 45 species, enabling robust alignments (Fig. S6A). These precursor miRNAs were distributed unevenly across the genome: chromosomes 14 and X contained the highest numbers (N = 14 each), while chromosome 6 contained none, and three miRNAs lacked corresponding human annotations in MirGeneDB (Fig. S6B). Of the 152 mature miRNAs, 80 originated from the 5’ arm and 72 from the 3’ arm (Fig. S6C).

**Figure 5:**
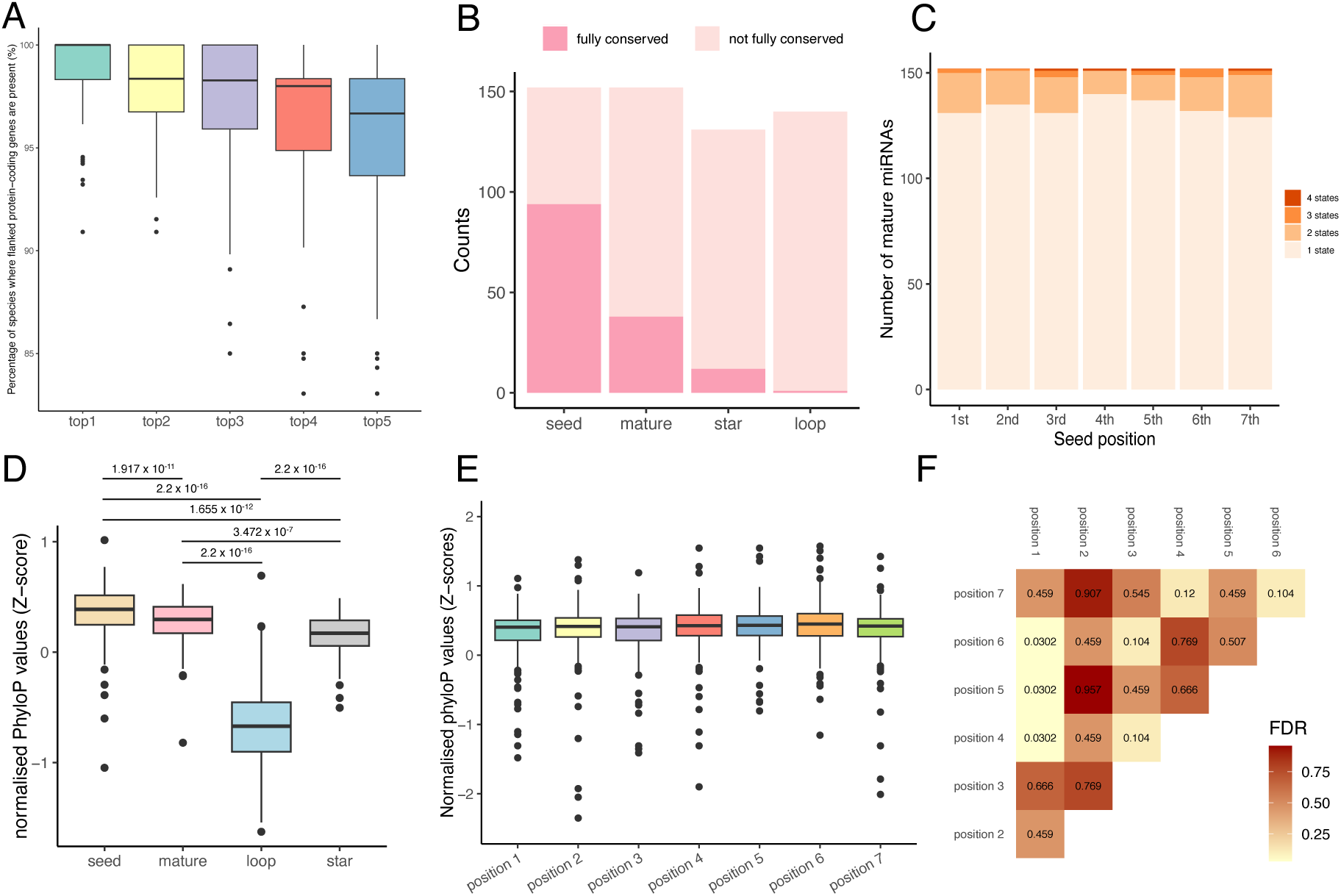
Analyses of orthologous miRNA families. **A)** Orthology validation of 143 individual precursor miRNAs using genome microsynteny. The boxplot shows the occurrence frequency of the top five protein-coding genes most consistently found within conserved syntenic blocks across species. **B)** Sequence conservation of orthologous miRNAs based on precursor alignments. Seed, mature, loop, and star regions were defined using human annotations from MirGeneDB as reference. **C)** Frequency of inferred states at each seed position based on precursor alignments of 143 orthologous miRNAs. “State 1” indicates a fully conserved nucleotide (one nucleotide type at the site), while “State 2, 3, and 4” indicate variability (two or more nucleotide types, e.g. A or T, G or deletion). **D)** Conservation levels of seed, mature, loop, and star regions estimated using phyloP scores (normalised z-scores) computed on precursor alignments of the 143 orthologous miRNAs. Statistical comparisons among regions were performed with Mann-Whitney *U* tests. **E)** Distribution of phyloP scores (normalised z-scores) for each seed position. **F)** Heatmap showing the FDR-adjusted p-values from pairwise Mann-Whitney *U* tests comparing phyloP score distributions among individual seed positions.

Across the 143 precursor miRNA alignments, 38 of 152 (25%) mature miRNAs were fully conserved in sequence, and 94 (61.8%) showed complete conservation of the seed region (Fig. 5B). Among the 131 star sequences, 12 (9.2%) star regions were fully conserved, and only a single loop region (miR-137-3p) was fully conserved (Fig. 5B). Examination of seed regions (position 2 to 8 of the mature miRNA) further showed that seed position 4 (corresponding to position 5 of the mature miRNA) exhibited the highest proportion of conserved homologous sites (92.1%; 1 state), followed by seed position 5 (90.1%), whereas seed position 7 showed the lowest proportion (84.9%) (Fig. 5C; Fig. S7). Next, we computed conservation (phyloP) scores for each homologous site across the 143 precursor miRNA alignments (see Methods). Seed regions exhibited the strongest conservation levels, followed by mature, star and loop regions, with all pairwise differences being statistically significant (P < 0.05; Mann-Whitney *U* test; Fig. 5D). Focusing specifically on individual seed positions, the first nucleotide of the seed (corresponding to position 2 of the mature miRNA) exhibited statistically significant lower conservation than seed positions 4, 5, and 6 (FDR < 0.05; Mann-Whitney *U* test; Figs. 5E-F). Seed positions 4, 5, and 6 also showed higher conservation levels than seed positions 2, 3, and 7, although these differences were not statistically significant (FDR > 0.05; Mann-Whitney *U* test; Figs. 5E-F).

### Ancestral state reconstruction of miRNA seed regions

Because the miRNA seed is the primary determinant of target recognition and regulatory function, we reconstructed the ancestral states of miRNA seed regions using the 143 precursor miRNA alignments and identified distinct evolutionary signatures in several families. Notably, miR-3660-3p and miR-493-5p exhibited clade-specific seed changes (Figs. 6A-B). For miR-3660-3p, position 4 of the seed was inferred to be adenine (A) at the mammalian ancestral node, but this site shifted to thymine (T) specifically within Chiroptera (Fig. 6A). miR-493-5p showed a different evolutionary pattern: seed position 7 was reconstructed as thymine (T) in the mammalian ancestor, whereas species belonging to Atlantogenata exhibited a derived substitution to cytosine (C) (Fig. 6B). Beyond these examples, several families displayed recurrent seed modifications across distantly related lineages (Figs. 6C-F). For instance, in miR-147-3p, seed position 6 repeatedly mutated to adenine (A) in marsupials, certain bat species, and *C. cristata*, whereas the remaining species retained guanine (G) at this position (Fig. 6C). Similarly, position 2 of the miR-676-3p seed was inferred to be guanine (G) in the mammalian ancestor, but recurrent adenine (A) substitutions arose independently in *C. didactylus*, *Tupaia chinensis*, *Otolemur garnettii*, *Rhinopithecus bieti*, and *S. syndactylus* (Fig. 6F). In addition, some families exhibited more complex, homoplasious evolutionary trajectories. For example, position 3 of the miR-3187-3p seed was reconstructed as cytosine (C) in the mammalian ancestral node, followed by a single C to G substitution in the ancestor of Haplorrhini and an independent C to A substitution in *Oryctolagus cuniculus* (Fig. 6H). Sequence alignments for all these cases are provided in Fig. S8.

**Figure 6:**
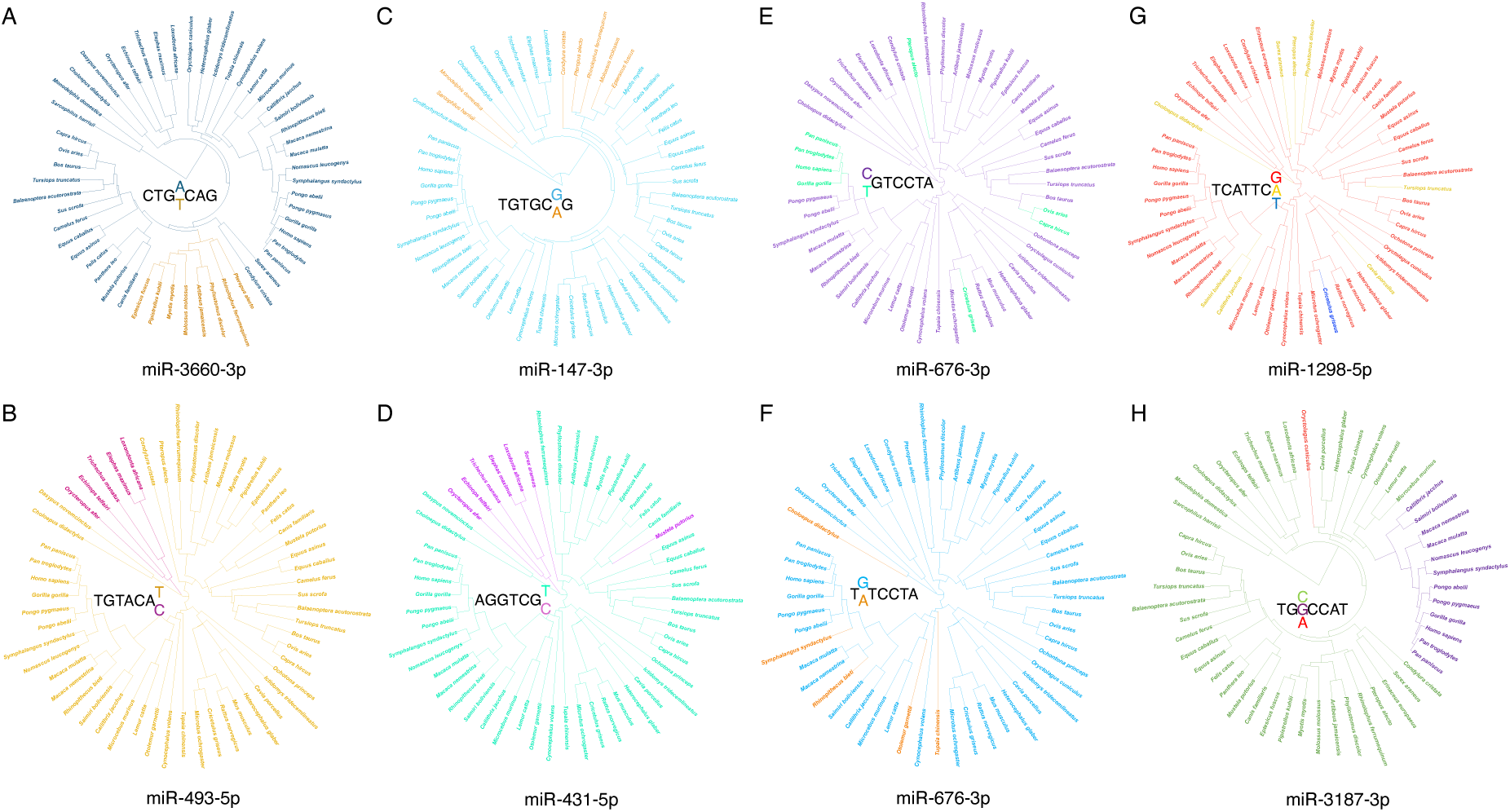
Examples of seed evolution across mammals. **A-B)** Instances of seed substitution that originated in ancestral lineages. **C-F)** Cases of recurrent seed mutations that evolved independently in distantly related clades. **G-H)** Examples illustrating homoplasious and more complex seed evolution. In each panel, the seed sequence is shown in the center, and sites exhibiting alternative mutations are highlighted in color.

## Discussion

Despite extensive study of protein-coding gene evolution, our understanding of miRNA diversification and evolutionary history across mammals remains limited. Much of this stems from the challenge of accurately and consistently annotating miRNA genes across diverse genomes, which complicates large-scale comparative studies (Tarver et al. 2018). Although tools such as Infernal and MirMachine enable genome-wide miRNA prediction independent of RNA-Seq, their inherent biases influence comparative analyses. Infernal often overpredicts due to inconsistently curated Rfam models, whereas MirMachine excels at conserved families but misses many divergent or lineage-specific miRNA families. This likely explains the large discrepancy in curated mammalian miRNA families between the two databases (Fig. 1A) and miRNA family predictions between Infernal and MirMachine (Fig. 1B).

In this study, we implemented a homology-based approach that mapped high-confidence mammalian miRNAs from MirGeneDB onto genomes to identify miRNA loci. Despite differences in the number of miRNA families predicted by each method, using consensus-based evaluation relative to multi-method agreement, our homology-based approach achieved a balanced performance, showing a better trade-off between sensitivity and specificity than either MirMachine or Infernal (Fig. 1C). While these consensus-based comparisons provided useful relative performance metrics, future validation against experimentally confirmed miRNA sets will be important to establish absolute accuracy rates. For families detected by all three approaches, copy number estimates from our method exhibited higher concordance with those from MirMachine than with Infernal (Fig. 1D). Moreover, among the 19 species with curated miRNA annotations in MirGeneDB, our approach recovered on average 10.7% more families (Fig. 2B). For example, MirGeneDB currently annotates MIR-6715 in 14 species, whereas our approach identified this family in 38 additional mammalian genomes, with the seed sequence fully conserved across all detections (Fig. S7). Likewise, MirGeneDB reports MIR-3660 only in cow, while our method recovered this family in 51 additional species and uncovered a clade-specific seed substitution in miR-3660-3p (Fig. 6A; Fig. S8). Furthermore, although MirMachine defines families largely based on shared seed sequences, we uncovered a plethora of miRNA families exhibiting seed alterations during evolution (Figs. 6A-H). Together, these findings indicate that our homology-based strategy enhances specificity by avoiding low-confidence CMs, while retaining sensitivity for moderately divergent family members that may not be captured by CM-based tools.

Application of the homology-based method to 73 mammalian species revealed that the largest bursts of miRNA family gains occurred at the nodes of Eutheria, Marsupialia, and Haplorrhini (Fig. 2A). This pattern is consistent with previous studies showing the early mammalian diversification was accompanied by accelerated miRNA birth, particularly within therian lineages where regulatory innovation expanded markedly (Christodoulou et al. 2010; Berezikov 2011). Similar miRNA expansions in Primates, especially Haplorrhines, have also been linked to lineage-specific regulatory innovations associated with neural and reproductive evolution (Noguer-Dance et al. 2010; Diaz et al. 2020). In contrast, the greatest miRNA losses were concentrated in Eulipotyphla and Scandentia (Fig. 2A). Elevated loss in these clades aligns with evidence that small, rapidly evolving mammalian lineages often exhibit high genomic turnover and reduced retention of newly emerged or weakly constrained miRNAs (Meunier et al. 2013), paralleling patterns of loss observed for other molecular markers in these groups (Uvizl et al. 2024). Both Eulipotyphlans and Scandentia exhibit elevated molecular evolutionary rates and signatures of dynamic genome evolution relative to other mammalian orders (Fan et al. 2013; Wu et al. 2017), conditions likely to promote increased genomic turnover and, consequently, miRNA loss. These results imply that miRNA innovation and decay are highly uneven across the mammalian tree, reflecting lineage-specific regulatory pressures that may have contributed to major evolutionary transitions.

While most miRNA families showed conserved copy numbers across mammals, reflected by high phylogenetic signals (Pagel’s λ) and low evolutionary rates (σ²), a subset (e.g., MIR-145, MIR-2285, MIR-544) displayed clear lineage-specific accelerations in duplication (Fig. 3B). One prominent example is MIR-2285, which exhibited notably high copy numbers within Cetartiodactyla, with no other mammalian lineages possessing this family (Fig. 3C). Although previously considered ruminant-specific (Bilbao-Arribas et al. 2023), we found that cetaceans (*Balaenoptera acutorostrata* and *Tursiops truncatus*) also carry high copy numbers, indicating that MIR-2285 originated and expanded earlier, likely in the common ancestor of Cetartiodactyla, rather than being restricted to terrestrial ruminants. In sheep, 146 MIR-2285 loci have been reported to express precursors across multiple tissues, with highest expression in immune-related organs such as lymph nodes (Bilbao-Arribas et al. 2023), a pattern also observed in other ruminants where MIR-2285 has been linked to regulation of inflammatory response and insulin resistance (Bao et al. 2013; Lawless et al. 2014). Many of the duplicated copies exhibit seed shifts or point mutations, suggesting potential diversification of regulatory targets. In our dataset, species within Cetartiodactyla carried an average of 1,219 copies, 99.8% of which overlapped with TEs (Fig. 3D). According to the previous findings, this highlights that TE content alone cannot reliably indicate whether TE-derived duplications have lost function without also considering secondary structure, expression evidence and additional functional metrics.

A key highlight of our study is the ability to resolve and confirm miRNA orthology relationships, even within paralog-rich families such as LET-7 (Fig. 4; Fig. 5A; Figs. S4-5). Because miRNAs are short and often conserved, sequence similarity alone rarely provides sufficient resolution for orthology assignment (Fromm et al. 2015; Langschied et al. 2023). By incorporating synteny context with homology-based predictions, we inferred orthologs from lineage-specific paralogs and accurately reconstructed miRNA family histories across mammals. Among 143 synteny-resolved miRNA orthologs, a large proportion showed high evolutionary conservation, as reflected by complete conservation of their mature or seed sequences (Fig. 5B), indicative of strong functional constraints. Our phyloP analyses further revealed a clear hierarchy of constraint across hairpin regions: the seed region exhibited the strongest conservation, followed by the mature sequence, the star sequence, and much lower conservation in the loop (Fig. 5D). This pattern reflects the central functional importance of the seed for target recognition and repression (Lewis et al. 2003). Changes in the seed region would likely disrupt regulatory interactions and are therefore strongly purged by purifying selection, while the non-seed portion of the mature miRNA and the star sequence retain moderate constraint because both can contribute to target pairing, stability, or occasional regulatory activity (Grimson et al. 2007). In contrast, the loop appears to evolve under relaxed constraint, accumulating mostly neutral substitutions without compromising hairpin folding or processing (Saunders et al. 2007; Bartel 2009). Collectively, these sharply stratified conservation patterns support the view that evolutionary conservation of miRNAs is tightly linked to functional necessity, with strong purifying selection acted on regions essential for regulatory function and relaxed constraints on structurally permissive regions.

Although the miRNA seed region is generally under strong constraint, we found that seed positions 4, 5, and 6 seem to be slightly more conserved than other seed positions (Figs. 5E-F). Previous studies have shown that mismatches in the centre of the seed region disrupt target recognition most severely because they interrupt continuous base-pairing and maximally destabilise the helix (Grimson et al. 2007; Bartel 2009). In contrast, although seed position 1 contributes to miRNA-mRNA interaction, evidence from studies in humans and *C. elegans* suggests that its role is more context dependent, as it is relatively permissive to non-Watson-Crick pairing (Grosswendt et al. 2014), and may contribute less to binding energetics than seed positions 4, 5, and 6 in canonical targeting interactions. These results suggest that the strength of purifying selection across the seed region is not uniform, with the strongest constraint likely acting on the center of the seed, reflecting their critical functional importance in miRNA-mRNA recognition. It is important to note, however, that our seed definition is based on human miRNA annotations from MirGeneDB. During evolution, some miRNAs undergo seed shifts (e.g., miR-337 in rodents (Clarke et al. 2025), not included in our analyses), switch their dominant mature arm (Kim et al. 2020), or become pseudogenised, all of which can complicate mature or seed recognition and interpretation in certain lineages. While such cases may introduce errors for a small subset of miRNAs, they are relatively uncommon, and thus the overall conservation pattern we describe should remain robust for the vast majority of miRNAs examined.

By aligning orthologous miRNAs across species, we also identified several cases in which miRNAs show distinct signatures of seed evolution (Figs. 6A-H). Seed mutation creates new targets (gain-of-function) and loses original targets (loss-of-function) (Bhattacharya and Cui 2017). These cases indicate that, despite the well-established functional constraint on the miRNA seed region, seed evolution in mammals is not strictly static. Instead, it reveals clade-specific, recurrent, or lineage-divergent patterns that point to a more nuanced evolutionary landscape. The clade-specific seed substitution observed in miR-3660-3p and miR-493-5p suggest that certain lineages may have experienced shifts in selective pressures or regulatory demands that favored modifications to particular seed positions, consistent with the arguments that miRNA evolution can contribute to lineage-specific innovation (Berezikov 2011). Notably, similar clade-specific seed change has been reported for miR-337-3p in bats (Jebb et al. 2020). Although specific biological consequences remain to be tested, such changes have the potential to alter the spectrum of miRNA targets, thereby contributing to lineage-specific regulatory rewiring. Similarly, families showing recurrent seed substitutions across phylogenetically distant species, such as miR-147-3p and miR-676-3p, may reflect instances of parallel evolution, where different lineages independently acquire similar seed changes in response to broadly comparable ecological or physiological pressures (Wang et al. 2024). In contrast, families exhibiting more complex and homoplasious trajectories, such as miR-3187-3p, highlight that seed evolution can take multiple independent paths even within closely related clades (Marco et al. 2010), suggesting that evolutionary constraints on the seed can be relaxed under certain lineage-specific contexts. Given that most genome assemblies analysed consist of a single haplotype and are not fully phased, it remains possible that some observed seed variants, especially the ones showing complex trajectories, reflect heterozygous positions rather than fixed substitutions. Although this is unlikely to influence the broader patterns we report, it is an important caveat when interpreting individual cases. Altogether, these findings suggest that while most miRNA seeds remain highly conserved, selective pressures acting at different evolutionary depths can drive modest but biologically meaningful divergence in miRNA targeting potential across species.

Elucidating miRNA evolution across mammals has long been challenged by the difficulty of accurately predicting miRNA families and, critically, establishing their orthology. In this study, we developed a homology-based framework to identify miRNA families and reconstruct their evolutionary histories across diverse mammalian genomes. While no single method can fully resolve all ambiguities inherent to miRNA annotation, our combined use of sequence similarity and genome syntenic context, together with stringent pipeline controls, provides a prudent framework that helps minimise false positives while preserving reasonable sensitivity. Independent lines of evidence from miRNA gain and loss patterns, conserved trajectories of copy number evolution, synteny-supported orthology assignments, conservation profiles of precursor components, and evolutionary signatures within seed regions collectively indicate that our predictions are broadly reliable, even if some individual cases warrant careful interpretation. Our findings suggest that although most miRNA families maintain stable copy numbers and seed identities over long evolutionary time, lineage-specific expansions and seed modifications do occur and have the potential to alter regulatory landscapes. Future work integrating small RNA-Seq data will be important for recovering novel and lineage-specific miRNA families. Taken together, our results provide a comprehensive genome-wide perspective on miRNA evolution, highlighting both deeply conserved features and more subtle, lineage-specific changes that may have contributed to the emergence of distinctive mammalian traits.

## Methods

### Concordance analysis of miRNA predictions from Infernal, MirMachine, and the homology-based approach

To evaluate the performance of our homology-based approach, we compared its miRNA predictions with those produced by Infernal and MirMachine using ten high-quality, phylogenetically representative mammalian genomes (*H. sapiens*, *M. musculus*, *Bos taurus*, *B. acutorostrata*, *Sus scrofa*, *Pipistrellus kuhlii*, *Loxodonta africana*, *Canis familiaris*, *S. harrisii*, and *O. anatinus*).

For the homology-based method, all mammalian precursor miRNA sequences were downloaded from MirGeneDB (v3.0) (Clarke et al. 2025), comprising 7,556 sequences from 19 mammalian species. Redundant sequences were removed using CD-HIT (v4.8.1) (Li and Godzik 2006) with a 90% similarity threshold, and excessively long sequences (>120bp) were removed, yielding a final query dataset of 1,774 precursor miRNAs representing 568 miRNA families (Table S1). Each genome was indexed using bowtie2-build (v2.5.2), and mapping was performed with Bowtie2 (v2.5.2) in end-to-end mode with increased sensitivity (-D 20 -R 3 -N 1 -L 20 -i S,1,0.50 --mp 3 -a) (Langmead and Salzberg 2012), reporting all valid alignments and allowing more divergent alignments to capture evolutionarily distant miRNA homologs. SAM files were sorted using Samtools (v1.19.2) (Li et al. 2009) and overlapping genomic hits arising from similar query sequences were collapsed to unique loci using Bedtools (v2.31.1) (Quinlan and Hall 2010). Notably, in the homology-based workflow, once a genomic locus was mapped by a MirGeneDB precursor sequence, it was counted as a copy of that miRNA family, regardless of seed conservation. This definition enables the detection of potential seed shifts or functional divergence within conserved precursor loci.

For comparative prediction, MirMachine (v0.2.13) (Umu et al. 2023) was run using the deuterostomia model with the --add-all-nodes option, and only high-confidence miRNA families defined at mammalian lineage nodes were retained. Infernal-based predictions were generated using Rfam miRNA covariate models (v15.0) annotated for mammals (Ontiveros-Palacios et al. 2025) and executed with cmsearch (v1.1.5) (Nawrocki and Eddy 2013), with --cut_ga flag to enforce family-specific gathering thresholds. Predictions with E-value < 10^-6^ were considered as high-confidence and retained.

To assess concordance among the three pipelines, we first manually ensured the consistency of miRNA family nomenclature between Rfam and MirGeneDB. Next, we evaluated the pseudo-sensitivity and pseudo-specificity for each method. Due to the absence of experimentally validated gold-standard miRNA datasets for these species, we adopted a consensus-based approach to evaluate the agreement of these three methods. Specifically, miRNA families predicted by at least two of the three tools were treated as consensus positives, whereas the ones predicted by only one method were treated as consensus negatives. For each method, pseudo-sensitivity was calculated as the proportion of consensus-positive families detected, and pseudo-specificity as the proportion of consensus-negative families correctly excluded. These metrics measure concordance with cross-method consensus rather than absolute biological accuracy. For miRNA families detected by all three pipelines, we quantified copy number consistency using Spearman’s rank correlation with the *cor.test* R function. We additionally examined miRNA families that were uniquely detected by the homology-based method across the ten species using the *UpSetR* (v1.4.0) R package (Conway et al. 2017).

### Homology-based miRNA prediction across 73 mammalian genomes

To investigate the evolutionary history of miRNA families across mammals, we applied the homology-based pipeline to 73 mammalian genomes, including the ten species used earlier for comparing the three prediction frameworks. These genomes represent all major mammalian lineages and encompass substantial ecological and morphological diversity, including Euarchontoglires (Primates, N = 21; Rodentia, N = 7; Lagomorpha, N = 2; Scandentia, N = 2; Dermoptera, N = 1), Laurasiatheria (Chiroptera, N = 8; Carnivora, N = 6; Cetartiodactyla, N = 7; Perissodactyla, N = 4; Eulipotyphla, N = 2), and Afrotheria (Afrosoricida, N = 1; Proboscidea, N = 2; Sirenia, N = 2, Tubulidentata, N = 1), with additional non-placental representatives from Monotremata (N = 1) and Marsupialia (Dasyuromorphia, Didelphimorphia, Diprotodontia, each N = 1), and Xenarthra (Cingulata, N = 1; Pilosa, N = 2). Notably, 54 out of the 73 species currently lack curated miRNA annotations in MirGeneDB (v3.0). When available, RefSeq genome assemblies were used, and for each genome, contigs shorter than 50 kb were excluded. Genome completeness was evaluated using BUSCO (v5.8.0) (Simao et al. 2015), with an average completeness of 94.4% ± 1.7% (Table S2). All genomes were soft masked using RepeatMasker (v4.1.8) (Smit 2013-2015) with default parameter settings prior to miRNA identification. miRNA prediction followed the same homology-based pipeline and filtering criteria described above, resulting in a copy-number matrix of 568 miRNA families across the 73 species (Table S3). Genome-wide miRNA coordinates (BED format) are available in Table S4.

### Gains and losses of miRNA families along the mammalian phylogenetic tree

Next, we inferred the gain and loss history of each miRNA family using maximum-likelihood ancestral state reconstruction implemented in *ape* R package (v5.8-1) (Paradis and Schliep 2019). Binary presence-absence states were fitted to a fixed, time-calibrated mammalian phylogeny obtained from TimeTree (v5) (Kumar et al. 2022) under a discrete equal-rates (ER) model, and ancestral states were assigned based on maximum posterior probability. Gains and losses were then mapped to individual branches by comparing inferred parent-child states (0 to 1 = gain; 1 to 0 = loss). A complete list of inferred gains and losses for all species is provided in Table S5.

### Analyses of miRNA family copy number

Principal component analysis (PCA) was performed using the *prcomp* function in R on log-transformed copy numbers of 568 miRNA families across 73 species. To explore the evolutionary dynamics of miRNA expansion, we quantified phylogenetic signals (Pagel’s λ) and evolutionary rates (σ^2^) for each family. Pagel’s λ measures the extent to which variation in copy number follows the phylogeny (λ = 0: no phylogenetic structure; λ = 1: variation fully consistent with phylogeny) (Pagel 1999), whereas σ^2^ estimates the rate of copy number evolution under a Brownian motion model, with larger σ^2^ values indicating faster change (Revell and Collar 2009). Analyses were performed using the same time-calibrated mammalian phylogeny described above. To investigate whether rapidly expanding families may include non-functional copies, we selected the 10 most variable families (highest σ^2^ values) and examined TE overlap in their precursor sequences. Copies without TE overlap were classified as high-confidence, whereas copies overlapping annotated TEs were considered as low-confidence. Copy number patterns of these families were further visualised across species.

### miRNA orthology assignments across species

Because miRNA sequences are short and often highly conserved, sequence similarity alone cannot reliably resolve orthology, particularly for multi-copy families. Therefore, we applied a synteny-based approach (Langschied et al. 2023; Uvizl et al. 2024) in which the relative positions of flanking protein-coding genes serve as genomic anchors for identifying orthologous miRNA loci across species. Of the 73 genomes examined, 61 with available gene annotations were used. miRNA loci identified from the homology-based pipeline were intersected with genome annotations using Bedtools (v2.31.1) (Quinlan and Hall 2010). For each miRNA locus, the five nearest upstream and five nearest downstream protein-coding genes were retrieved to define a synteny block. These blocks were compared across species to determine whether miRNAs were located within homologous genomic neighborhoods. This approach was applied to resolve orthology for single-copy, two-copy miRNA families, and for the multi-copy LET-7 family. Orthology confidence was quantified by calculating, for each synteny block, the frequency at which the top five most recurrent anchor genes appeared across species.

### Evolutionary Analyses of Orthologous miRNAs

To ensure robust comparative analysis, only synteny-supported orthologous miRNAs present in more than 45 of the 61 annotated genomes (>75% species representation) were included. We excluded the LET-7 family from this analysis to focus on single- and two-copy miRNA families, as its extreme conservation, functional redundancy, and large number of paralogous loci would disproportionately influence conservation estimates and complicate interpretation due to non-independence among copies. Precursor sequences were aligned using MUSCLE (v3.8.425) (Edgar 2004) in AliView (v1.28) (Larsson 2014), and alignments were manually curated to remove large insertions or poorly aligned regions. Site-wise conservation was computed using the phyloP module of the Phylogenetic Analysis with Space/Time Models (PHAST) package (Pollard et al. 2010), guided by the mammalian time-scaled phylogeny aforementioned. Species lacking a given miRNA were pruned from the tree using *ete3* (v3.1.3) in Python (v3.11.7) (Huerta-Cepas et al. 2016). For each alignment, a neutral substitution model was first estimated using phyloFit under the HKY85 model, and phyloP was run in CONACC mode to detect homologous sites showing conservation or acceleration relative to the neutral expectation across the precursor miRNA hairpin.

To allow direct comparison among miRNAs, site-specific phyloP scores were normalised to z-scores. The seed, mature, star and loop regions were defined according to MirGeneDB (v3.0), using the human annotation as reference. For each alignment, the mean of z-score-normalised phyloP values was calculated for the seed, mature, star and loop regions, respectively. Note that some miRNAs have a mature and co-mature sequence defined but no annotated star sequence (e.g., miR-28), and miR-451 lacks a defined star sequence entirely. Evolutionary change within the seed was quantified by counting the number of unique states observed at each seed position (bases 2-8 of the mature strand). A value of 1 state indicates complete conservation, whereas 2 - 4 states indicate the presence of substitutions and/or deletions at that homologous site. Multiple sequence alignments of synteny-supported orthologous miRNAs included in this analysis are provided on GitHub (https://github.com/sarahjanepower/homology).

### Ancestral state reconstruction of miRNA seed regions

To investigate how seed sequences evolved across mammals, we reconstructed the ancestral state of each seed position for orthologous miRNAs using PastML (v1.9.50) (Ishikawa et al. 2019). PastML applies a probabilistic parsimony framework, in this case the MPPA model (maximum parsimony with posterior probability approximation), to infer the most likely state (nucleotide or deletion) at each internal node of a phylogeny. Before running PastML, we removed seed alignment sites that showed no variation (i.e., only one state across all species), and treated each distinct nucleotide or deletion observed at a seed position as a separate discrete trait. The mammalian time-scaled phylogenetic tree described above served as the guide tree for inference. The inferred evolutionary transitions were plotted on the phylogeny using the iTOL web interface (v7.2.2) (Letunic and Bork 2021) to highlight lineage-specific seed evolution.

### Statistical analyses

All statistical analyses were performed in R (v4.5.1) (Team 2010). Spearman’s rank correlation, Mann-Whitney *U* tests, phylogenetic signal (Pagel’s λ) tests, evolutionary rate (σ^2^) analyses, and phylogenetic tree manipulations were conducted in R using standard functions and relevant packages. Where multiple tests were performed, P-values were corrected using the false discovery rate (FDR) method. Unless stated otherwise, results with FDR-adjusted P < 0.05 were considered statistically significant.

## Data Access

Information on all genomes analysed in this study is provided in Table S2. Multiple sequence alignments of orthologous precursor miRNAs, together with all scripts and code used for data analysis, are available on GitHub (https://github.com/sarahjanepower/homology).

## Acknowledgements

The work was supported by the Science Foundation Ireland (SFI) Centre for Research Training in Genomics Data Science (18/CRT/6214) and an Irish Research Council Laureate Award (IRCLA/2022/3212). We thank Bastian Fromm (Arctic University of Norway) and Sébastien J. Puechmaille (University of Montpellier) for valuable discussions and advice on data analysis and interpretation. We also acknowledge the UCD Sonic High Performance Computing facility for providing computational resources and support.

## Author contributions

Conceptualization: Z.H.; methodology: S.P., and Z.H.; software: S.P., and Z.H.; investigation: S.P., and Z.H.; writing: Z.H., and S.J.; funding acquisition: S.J., and Z.H.

